# Bagaza virus in wild birds, Portugal, 2021

**DOI:** 10.1101/2021.11.26.470055

**Authors:** João Queirós, Sílvia Barros, Alberto Sánchez-Cano, Margarida Henriques, Teresa Fagulha, Fábio Abade dos Santos, Margarida Duarte, Catarina Gonçalves, David Gonçalves, Marisa Rodrigues, Teresa Cardona Cabrera, Isabel G. Fernández de Mera, Christian Gortazar, Ursula Höfle, Paulo Célio Alves

## Abstract

Bagaza virus (BAGV) emerged in Spain, in 2010. Since then, BAGV was not reported in other European countries until September 2021, when BAGV was diagnosed by molecular methods in one corn bunting and several red-legged partridges, after abnormal mortality in Southern Portugal. Sequencing revealed high similarity with strains reported previously.

## Introduction

Bagaza virus (BAGV) is a positive-sense and single stranded RNA virus. It belongs to the mosquito-borne cluster of the genus *Flavivirus*, family *Flaviviridae* from the Ntaya serocomplex, and has a great nucleotide similarity with Israel turkey meningoencephalitis virus (ITV) (1). *Flavivirus* genus includes a number of pathogenic viruses associated with neurological disease in various wild and domestic animals, and in humans, such as Bagaza, Usutu, West Nile, Japanese Encephalitis (JEV), Dengue, Zika and Yellow Fever viruses (1). BAGV was first isolated in 1966 from a pool of *Culex* mosquitoes in the Bagaza district of Central African Republic. Subsequently, it was detected in several species of mosquitoes, more recently, in *Cx. perexiguus* in United Arab Emirates (9) and *Cx. univittatus* in Namibia (10). In vertebrates, BAGV associated-deaths were first detected in red-legged partridges (*Alectoris rufa*) and ring-necked pheasants (*Phasianus colchicu*) in Spain in 2010 (2), and a few years later, in 2016, in Himalayan monal (*Lophophorus impejanus*) in South Africa (6). BAGV infection causes neurological disease in red-legged partridges, grey partridges (*Perdix perdix*), ring-necked pheasants, and, to a lower degree, in common wood pigeons (*Columba palumbus*) (1,2-6). The estimated mortality rates range from 23% to 30% in naturally and experimentally infected red-legged partridges (4,7), reaching 40 % in experimentally infected grey partridges (5). The rates are lower in pheasants and in columbiformes (7,3). A study conducted in India in 1996, comprising 53 humans undergoing acute encephalitis showed a 15% positivity for BAGV neutralizing antibodies, suggesting its zoonotic potential (8). Despite sharing land borders with Spain, no evidence of BAGV circulation had been reported in Portugal. Here, we describe for the first time a BAGV outbreak associated with abnormal fatalities in red-legged partridges and one corn bunting (*Emberiza calandra*) in southern Portugal in autumn 2021.

### The study

On September 1^st^, 2021, three red-legged partridges were found dead in a hunting ground in Serpa, southern Portugal. One of them was in advanced stage of decomposition and was not analysed. Subsequently, ten wild birds (nine partridges and one corn bunting) were found dead between September and mid-October in the same area (Table 1). The sighting reports of partridges with compatible neurological signs such as disorientation and motor incoordination were also reported locally (*specimens not collected*). Laboratory examinations were initially carried out at the IREC and CIBIO-InBIO. Official diagnosis was carried later, in October 2021 at the National Reference Laboratory of Animal diseases of Portugal (INIAV, I.P.).

**Table 1:** Data on the BAGV-positive specimens analysed in this study.

| Code | Species | Status | Collection date | Sex | Age class | Real time RT-PCR (RT-qPCR) |  |  |  |  |  |  |  | Pan-Flavivirus conventional RT-PCR |  |
| --- | --- | --- | --- | --- | --- | --- | --- | --- | --- | --- | --- | --- | --- | --- | --- |
|  |  |  |  |  |  | Duplex RT-qPCR<sup>1</sup> (3' NTR) |  |  |  | Uniplex RT-qPCR<sup>2</sup> (NS5) |  |  |  | nested RT-PCR<sup>3</sup> (NS5) | RT-PCR<sup>4</sup> (NS2b) |
|  |  |  |  |  |  | Feather | Brain | Kidney | Spleen | Heart | Kidney | Brain | Intestine | Feather/Spleen | Heart |
| Partridge 1 | *A. rufa* | Found dead | 01/09/21 | Male | Adult | + | - | - | - | nt | nt | nt | nt | +(#) | nt |
| Partridge 2 | *A. rufa* | Found dead | 01/09/21 | Male | Adult | - | - | - | + | nt | nt | nt | nt | +(#) | nt |
| Partridge 3 | *A. rufa* | Found dead | 28/09/21 | Female | Juvenile | + | + | - | nt | nt | nt | nt | nt | - | nt |
| Partridge 4 | *A. rufa* | Found dead | 29/09/21 | Female | Juvenile | + | + | - | nt | nt | nt | nt | nt | +(#) | nt |
| Corn Bunting | *E. calandra* | Found dead | 09/10/21 | unkown | unkown | - | - | - | +(#) | nt | nt | nt | nt | - | nt |
| Partridge 7 | *A. rufa* | Found dead | 13/10/21 | Male | Adult | nt | nt | nt | nt | + | + | + | + | nt | +(#) |
| Partridge 8 | *A. rufa* | Found dead | 13/10/21 | Male | Adult | nt | nt | nt | nt | + | + | + | + | nt | +(#) |
| Partridge 9 | *A. rufa* | Found dead | 13/10/21 | Female | Juvenile | +(#) | +(#) | + | nt | nt | nt | nt | nt | +(#) | nt |
| Partridge 11 | *A. rufa* | Found dead | 14/10/21 | Female | Juvenile | + | + | + | nt | nt | nt | nt | nt | +(#) | nt |
| Partridge 14 | *A. rufa* | Live-trapped | 03/10/21 | Male | Adult | + | na | na | na | na | na | na | na | +(#) | na |
| Partridge 16 | *A. rufa* | Live-trapped | 03/10/21 | Male | Adult | + | na | na | na | na | na | na | na | - | na |
| Partridge 33 | *A. rufa* | Live-trapped | 03/10/21 | Male | Adult | + | na | na | na | na | na | na | na | - | na |
| Partridge 37 | *A. rufa* | Live-trapped | 03/10/21 | Male | Juvenile | + | na | na | na | na | na | na | na | - | na |

+, positive; -, negative; nt, not tested; na, not applicable; # - samples for which successful sequences were obtained. 1- dRT-qPCR developed by Lizalde et al., 2020 for the detection of JEV and virus from the Ntaya serocomplex; 2- RT-qPCR developed by Buitrago et al., 2012 for the specific detection of BAGV, 3- nested RT-PCR developed by Sánchez-Seco et al. (2005) for the detection of *Flavivirus*; 4- RT-PCR, developed by the INIAV team (unpublished)

**Table 2:** Information on the PCR methods used in this study for the molecular diagnosed of BAGV.

| System | Target region | Amplicon |  | Reference of the method |
| --- | --- | --- | --- | --- |
|  |  | Size | Position in sequence HQ644143 |  |
| dRT-qPCR | 3'NTR | 171 bp (108 nt)* | 10510 to 10680 | 11 |
| RT-qPCR | NS5 | 61 bp (ns) | 9056 to 9116 | 12 |
| Nested RT-PCR | NS5 | 143 bp (110 nt)* | 8996 to 9139 | 13 |
| In house RT-PCR | NS2b | 209 bp (171 nt)* | 5426 to 5634 | *Not published* |

\* Size of BAGV sequence without the primers' annealing sequences, used in BLAST and phylogenetic analyses; ns, not sequenced

Overall, 12 birds found dead were subjected to detailed necropsy during which samples from several organs were collected for further analyses. For histopathology, organs were fixed in 10% neutral buffered formalin. In addition, growing feathers were collected from 30 partridges live-trapped in the same area on October 3^rd^, using cages with wheat bait (Permission number 4 – DGVF/DRCA/ 2021). Viral RNA was extracted from samples of feather pulp, brain, heart, kidney, spleen and/or intestine using TRIzol reagent (Merck, Madrid, Spain), or the IndiMag Pathogen Kit (Indical, Leipzig, Germany) in a King Fisher Flex extractor (ThermoScientific, Waltham, EUA), following manufacturers’ instructions. Molecular detection was carried out at the three laboratories involved in testing (IREC, CIBIO-InBIO and INIAV), following two strategies targeting different regions of the BAGV genome (NS2b, NS5 and 3’NTR) (Table 1): *i*) a duplex RT-qPCR (dRT-qPCR) for the simultaneous and differential detection of JEV and Ntaya flavivirus serocomplexes, using the a method previously described (11), and *ii*) a uniplex RT-qPCR specific for NS5 coding region of BAGV (12). Subsequently, two pan-flavivirus conventional RT-PCRs were used, namely a nested RT-PCR previously described (13) and an *in house* RT-PCR (INIAV, *unpublished*) targeting part of the NS5 and NS2b genes, respectively. The amplicons were purified and directly sequenced using the ABI Prism BigDye Terminator v3.1 Cycle sequencing kit on a 3130 Genetic Analyser (Applied Biosystems, Foster City, CA, U.S.A.). These sequences were used for BLAST and phylogenetic analyses.

The results showed that eight red-legged partridges and one corn bunting out of the 12 wild birds found dead (66.7%), and four out of 30 live-captured red-legged partridges (13.3%) tested positive for BAGV (Table 1, Table S1), confirming the cause of death. Three 108 bp sequences obtained from dRT-qPCR from two dead birds (Partridge 9 and the bunting) showed 100% similarity in the 3’NTR region with BAGV genome strains published from the 2010 outbreak in Spain (HQ644143) (Table 1). Six 110 bp sequences obtained in nested RT-PCR from six partridges showed 99.1% similarity with the BAGV strains mentioned above, due to one mutation (T to C, position 9095 in HQ644143) in the NS5 region. Two 171 bp sequences obtained in RT-PCR from Partridges 7 and 8 showed 98.8% similarity with sequence HQ644143.

Two phylogenetic trees were constructed using Maximum Likelihood method and Kimura 2-parameter model, operating MEGA software. These were based on concatenated NS5/3’NTR (108 bp+110 bp; Fig 1a) and NS2b (171 bp; Fig1b) sequences retrieved from public databases. In both trees, the BAGV-Portugal 2021 sequences obtained from Partridge 9 (Fig 1a) and Partridge 7 (Fig 1b) grouped close to a cluster comprising strains from Spain-2010, Zambia-2013 and Namibia-2018.

**Figure 1.**
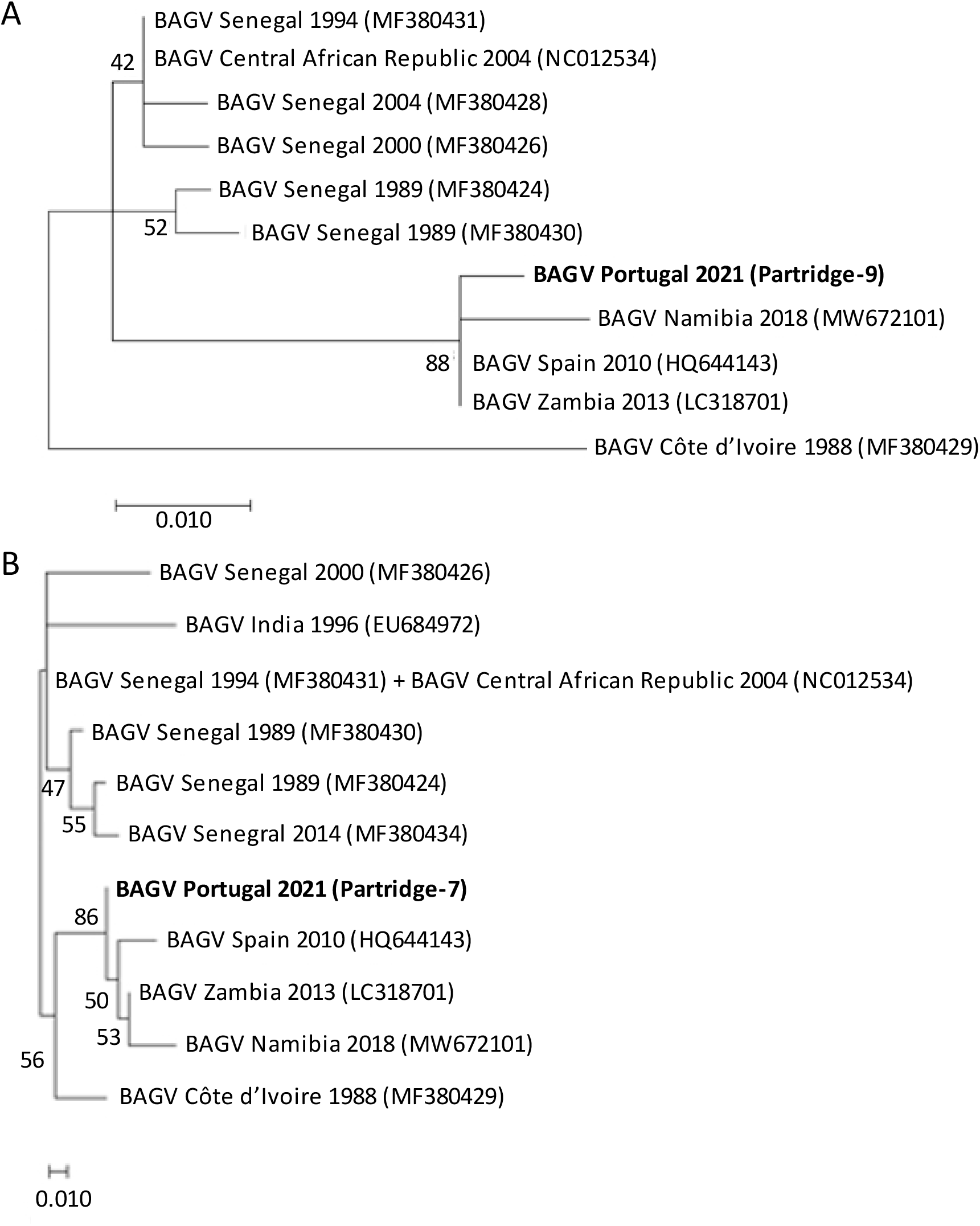
Maximum Likelihood trees based on concatenated 218bp of NS5/3’NTR (A) and 171bp of NS2b (B) from BAGV. Each of the two analyses involved 19 BAGV nucleotide sequences, with only one difference between trees, namely sequence EU684972, which is absent from Fig 1a, due to lack of 3’NTR region and, sequence MF380428 which is absent from Fig 1b, since it is identical to sequence MF380431. Duplicate sequences were removed from the alignments. Sequence HQ644143 is identical to HQ644144, KR108244, KR1082445 and KR108246; sequence MF380431 is identical to MF380425, MF380427, MF380432 and MF380433.The trees with the highest log likelihood are shown. The percentage of trees in which the associated taxa clustered together is shown next to the branches. Initial tree(s) for the heuristic search were obtained automatically by applying Neighbor-Join and BioNJ algorithms to a matrix of pairwise distances estimated using the Maximum Composite Likelihood (MCL) approach, and then selecting the topology with superior log likelihood value. A discrete Gamma distribution was used to model evolutionary rate differences among sites (5 categories (+G, parameter = 0.3621)). The trees are drawn to scale, with branch lengths measured in the number of substitutions per site. Codon positions included were 1st+2nd+3rd+Noncoding. Evolutionary analyses were conducted in MEGA X.

Upon necropsy, all birds were in good body condition suggesting an acute course of disease. Histopathology albeit hampered by autolysis and freezing artefacts revealed lymphoid depletion in the spleen, severe congestion and moderate to abundant diffuse mononuclear inflammatory infiltrates and focal necrosis in all tissues. The heart, brain, kidney and liver were the most affected organs (Fig 2).

**Figure 2:**
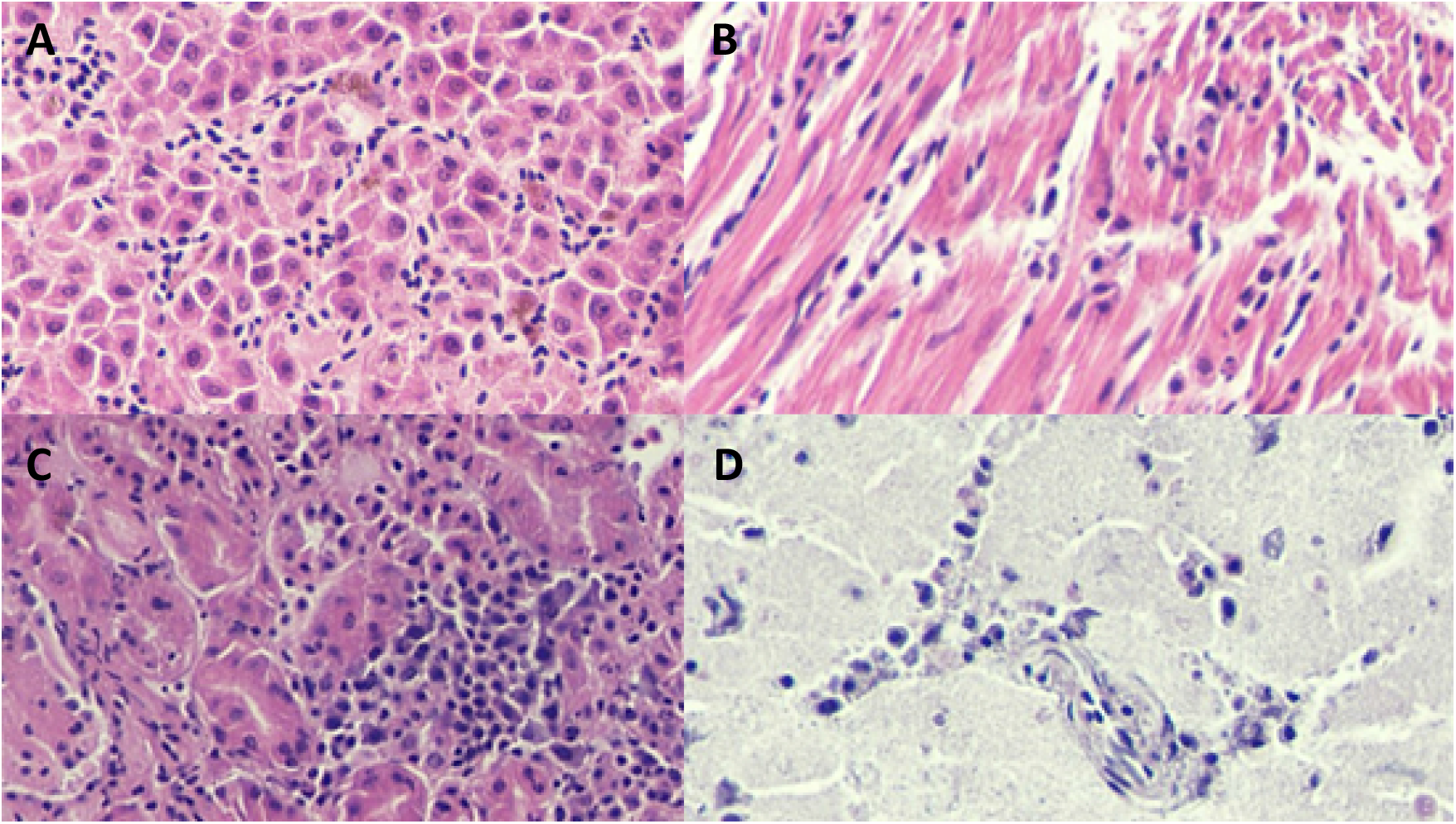
Microscopic lesions due to BAGV infection in red-legged partridges. Haematoxilin Eosin staining 400x magnification: A) liver: congestion, hemozoin presence in Kupffer cells, focal hepatocyte necrosis and a moderate mononuclear infiltrate are visible despite some freezing artefacts; B) heart: congestion, haemorrhage, oedema and degeneration of myofibers of the myocardium in addition to endothelial swelling and moderate to abundant diffuse mononuclear infiltrates; C) kidney: tubulointerstitial nephritis characterized by congestion, haemorrhage necrosis of proximal convoluted tubular epithelium and diffuse moderate to abundant mononuclear inflammatory infiltrate; D) brain: mild nonpurulent encephalitis with congestion mononuclear cell extravasation and endothelial cell swelling.

## Discussion and Conclusions

BAGV was detected for the first time in resident wild birds (red-legged partridges and corn bunting) in southern Portugal between September and October 2021. Moreover, 13.3% of live-captured red-legged partridges carried the virus in growing feathers, suggesting active circulation of BAGV in the region, evidencing the utility of growing feathers for BAGV detection in live birds in field studies. Red-legged partridge ecological and demographic studies are required to evaluate the extent and magnitude of BAGV outbreak in Portugal. However, significant population declines are expected based on mortality rate previously estimated for this species in the past (4,7). The additional finding of a fatal case in a songbird, the corn bunting, suggests that BAGV might have a broader spectrum and impact in wild bird species. An integrated wildlife monitoring (IVM) on a wide range of bird species is proposed to disclose the magnitude of BAGV spread in the wild (14). In addition, efforts are needed to identify the vectors for BAGV in Portugal and their role on the epidemiology of disease. *Culex univittatus* is one of the putative competent vectors (10) detected by the nationwide surveillance of haematophagous arthropods in Portugal (REVIVE). Despite the high similarity between the strains from the 2021 outbreak in Portugal and those from Cádiz 2010 (Southern Spain), given the small size of the sequences produced, further genomic analysis is needed to elucidate the phylogenetic relationships with previously known BAGV strains. No conclusions can yet be taken regarding the origin of the infection. However, the introduction by infected birds from North Africa or from Spain, or assuming that the disease may have persisted in southern Spain (15) constitute possible explanations.

## Acknowledgments

We thank personnel from the hunting sector for their help during field work (sampling and capturing), and Dr Fernanda Simões (INIAV) for the sex determination of Patridges 7 and 8 by molecular methods.

