## Supplementary material for "Bagaza virus in wild birds, Portugal, 2021": Table S1

Table S1: Data on the specimens analysed in this study relative to 40 Red-legged partridges (*Alectoris rufa*) and one corn bunting (*Emberiza calandra)*.

|  |  |  |  |  |  | **Real time RT-PCR (RT-qPCR)** | | | | | | | | |  | **Pan-Flavivirus conventional RT-PCR** | | |
| --- | --- | --- | --- | --- | --- | --- | --- | --- | --- | --- | --- | --- | --- | --- | --- | --- | --- | --- |
| **Code** | **Species** | **Status** | **Collection Date** | **Sex** | **Age class** | **Duplex RT-qPCR^1^**  **(3’NTR)** | | | |  | **Uniplex RT-qPCR^2^**  **(NS5)** | | | |  | **nested RT-PCR^3^**  **(NS5)** |  | **RT-PCR^4^**  **(NS2b)** |
|  |  |  |  |  |  | **Feathers** | **Brain** | **Kidney** | **Spleen** |  | **Heart** | **Kidney** | **Brain** | **Intestine** |  | **Feather/Spleen** |  | **Heart** |
| Partridge 1 | *A. rufa* | Found dead | 01/09/21 | Male | Adult | 22.77* | - | - | - |  | nt | nt | nt | nt |  | +(#) |  | nt |
| Partridge 2 | *A. rufa* | Found dead | 01/09/21 | Male | Adult | - | - | - | 26.82* |  | nt | nt | nt | nt |  | +(#) |  | nt |
| Partridge 3 | *A. rufa* | Found dead | 28/09/21 | Female | Juvenile | 26.76* | 32.18* | - | nt |  | nt | nt | nt | nt |  | - |  | nt |
| Partridge 4 | *A. rufa* | Found dead | 29/09/21 | Female | Juvenile | 29.72* | 37.87* | - | nt |  | nt | nt | nt | nt |  | +(#) |  | nt |
| Partridge 5 | *A. rufa* | Found dead | 04/10/21 | Male | Adult | - | - | - | nt |  | nt | nt | nt | nt |  | na |  | nt |
| Partridge 6 | *A. rufa* | Found dead | 08/10/21 | Female | Adult | - | - | - | nt |  | nt | nt | nt | nt |  | na |  | nt |
| Bunting 1 | *E. calandra* | Found dead | 09/10/21 | unkown | unkown | - | - | - | 43.27*(#) |  | nt | nt | nt | nt |  | - |  | nt |
| Partridge 7 | *A. rufa* | Found dead | 13/10/21 | Male | Adult | nt | nt | nt | nt |  | 20.40* | 24.50* | 21.52* | 36.71* |  | nt |  | +(#) |
| Partridge 8 | *A. rufa* | Found dead | 13/10/21 | Male | Adult | nt | nt | nt | nt |  | 31.71* | 34.16* | 29.06* | 39.36* |  | nt |  | +(#) |
| Partridge 9 | *A. rufa* | Found dead | 13/10/21 | Female | Juvenile | 26.41*(#) | 28.19*(#) | 31.98* | nt |  | nt | nt | nt | nt |  | +(#) |  | nt |
| Partridge 10 | *A. rufa* | Found dead | 14/10/21 | Female | Juvenile | - | - | - | nt |  | nt | nt | nt | nt |  | na |  | nt |
| Partridge 11 | *A. rufa* | Found dead | 14/10/21 | Female | Juvenile | 17.21* | 23.24* | 23.47* | nt |  | nt | nt | nt | nt |  | +(#) |  | nt |
| Partridge 12 | *A. rufa* | Live-trapped | 03/10/21 | Male | Juvenile | - | na | na | na |  | na | na | na | na |  | na |  | na |
| Partridge 13 | *A. rufa* | Live-trapped | 03/10/21 | Female | Juvenile | - | na | na | na |  | na | na | na | na |  | na |  | na |
| Partridge 14 | *A. rufa* | Live-trapped | 03/10/21 | Male | Adult | 35.12* | na | na | na |  | na | na | na | na |  | +(#) |  | na |
| Partridge 15 | *A. rufa* | Live-trapped | 03/10/21 | Female | Juvenile | - | na | na | na |  | na | na | na | na |  | na |  | na |
| Partridge 16 | *A. rufa* | Live-trapped | 03/10/21 | Male | Adult | 33.14* | na | na | na |  | na | na | na | na |  | - |  | na |
| Partridge 17 | *A. rufa* | Live-trapped | 03/10/21 | Female | Adult | - | na | na | na |  | na | na | na | na |  | na |  | na |
| Partridge 18 | *A. rufa* | Live-trapped | 03/10/21 | Female | Juvenile | - | na | na | na |  | na | na | na | na |  | na |  | na |
| Partridge 19 | *A. rufa* | Live-trapped | 03/10/21 | Male | Juvenile | - | na | na | na |  | na | na | na | na |  | na |  | na |
| Partridge 20 | *A. rufa* | Live-trapped | 03/10/21 | Male | Adult | - | na | na | na |  | na | na | na | na |  | na |  | na |
| Partridge 21 | *A. rufa* | Live-trapped | 03/10/21 | Female | Juvenile | - | na | na | na |  | na | na | na | na |  | na |  | na |
| Partridge 22 | *A. rufa* | Live-trapped | 03/10/21 | Male | Adult | - | na | na | na |  | na | na | na | na |  | na |  | na |
| Partridge 23 | *A. rufa* | Live-trapped | 03/10/21 | Male | Juvenile | - | na | na | na |  | na | na | na | na |  | na |  | na |
| Partridge 24 | *A. rufa* | Live-trapped | 03/10/21 | Male | Adult | - | na | na | na |  | na | na | na | na |  | na |  | na |
| Partridge 25 | *A. rufa* | Live-trapped | 03/10/21 | Male | Adult | - | na | na | na |  | na | na | na | na |  | na |  | na |
| Partridge 26 | *A. rufa* | Live-trapped | 03/10/21 | Female | Adult | - | na | na | na |  | na | na | na | na |  | na |  | na |
| Partridge 27 | *A. rufa* | Live-trapped | 03/10/21 | Female | Juvenile | - | na | na | na |  | na | na | na | na |  | na |  | na |
| Partridge 28 | *A. rufa* | Live-trapped | 03/10/21 | Male | Juvenile | - | na | na | na |  | na | na | na | na |  | na |  | na |
| Partridge 29 | *A. rufa* | Live-trapped | 03/10/21 | Male | Juvenile | - | na | na | na |  | na | na | na | na |  | na |  | na |
| Partridge 30 | *A. rufa* | Live-trapped | 03/10/21 | Male | Adult | - | na | na | na |  | na | na | na | na |  | na |  | na |
| Partridge 31 | *A. rufa* | Live-trapped | 03/10/21 | Female | Adult | - | na | na | na |  | na | na | na | na |  | na |  | na |
| Partridge 32 | *A. rufa* | Live-trapped | 03/10/21 | Male | Juvenile | - | na | na | na |  | na | na | na | na |  | na |  | na |
| Partridge 33 | *A. rufa* | Live-trapped | 03/10/21 | Male | Adult | 34.42* | na | na | na |  | na | na | na | na |  | - |  | na |
| Partridge 34 | *A. rufa* | Live-trapped | 03/10/21 | Male | Adult | - | na | na | na |  | na | na | na | na |  | na |  | na |
| Partridge 35 | *A. rufa* | Live-trapped | 03/10/21 | Female | Juvenile | - | na | na | na |  | na | na | na | na |  | na |  | na |
| Partridge 36 | *A. rufa* | Live-trapped | 03/10/21 | Female | Juvenile | - | na | na | na |  | na | na | na | na |  | na |  | na |
| Partridge 37 | *A. rufa* | Live-trapped | 03/10/21 | Male | Juvenile | 35.05* | na | na | na |  | na | na | na | na |  | - |  | na |
| Partridge 38 | *A. rufa* | Live-trapped | 03/10/21 | Male | Juvenile | - | na | na | na |  | na | na | na | na |  | na |  | na |
| Partridge 39 | *A. rufa* | Live-trapped | 03/10/21 | Female | Juvenile | - | na | na | na |  | na | na | na | na |  | na |  | na |
| Partridge 40 | *A. rufa* | Live-trapped | 03/10/21 | Male | Juvenile | - | na | na | na |  | na | na | na | na |  | na |  | na |
| Partridge 41 | *A. rufa* | Live-trapped | 03/10/21 | Female | Juvenile | - | na | na | na |  | na | na | na | na |  | na |  | na |

+, positive; -, negative; nt, not tested; na, not applicable; # - samples for which successful sequences were obtained.1- dRT-qPCR developed by Lizalde et al., 2020 for the detection of JEV and virus from the Ntaya serocomplex; 2- RT-qPCR developed by Buitrago et al., 2012 for the specific detection of BAGV (*-Cq values),3- nested RT-PCR developed by Sánchez-Seco et al. (2005) for the detection of Flavivirus; 4- RT-PCR, developed by the INIAV team (unpublished)
